# Voronoi tessellation as a complement or replacement for confidence ellipses in the visualization of data projection and clustering results

**DOI:** 10.1101/2025.09.18.677103

**Authors:** Jörn Lötsch, Dario Kringel

**Author notes:** **Correspondence to:** Prof. Dr. Dr. Jörn Lötsch, Goethe - University, Theodor - Stern - Kai 7, 60590 Frankfurt am Main, Germany, E-Mail.

## Abstract

Visualizing two-dimensional data projections with group-wise coloring and confidence ellipses is a standard approach in biomedical data analysis. However, this method can obscure subtle group overlaps or atypical cases. Voronoi tessellation, which is widely used in crystallography to analyze local structure, offers a parameter-free geometric alternative that may improve the evaluation of group structure in projected data. We compared the use of confidence ellipses and Voronoi tessellation as visualization tools for two-dimensional data projections on three artificial datasets and three biomedical datasets. For datasets with well-separated classes, both visualization techniques effectively delineated groups. Voronoi tessellation more clearly highlighted cases with projections overlapping the opposite group and revealed internal group heterogeneity. In datasets with moderate or absent group separation, Voronoi tessellation more effectively exposed the lack of meaningful structure. Meanwhile, confidence ellipses more clearly indicated distant outliers. Voronoi tessellation also facilitated the identification of clustering failures. Thus, Voronoi tessellation enhances the detection of deviations from expected group patterns and provides geometric insights that complement statistical summaries from confidence ellipses. Therefore, integrating Voronoi tessellation into standard data analysis workflows is a valuable addition for visualizing projected biomedical data and supports hypothesis validation and exploratory analyses in dimensionality reduction and clustering.

## Introduction

Projecting data and inspecting group separation are standard parts of biomedical data analysis workflows. Unsupervised or supervised techniques, such as principal component analysis (PCA [1, 2]) or partial least squares discriminant analysis (PLS-DA [3]), respectively, are among commonly used methods to investigate study group separation, with many further methodological alternatives [4]. Consequently, standard software implementations, whether in data science-oriented languages such as R [5] or Python [6], as well as commercial biomedical data analysis packages, offer data projections onto the two-dimensional ℝ^2^ plane with group-wise coloring of points. The projection of subgroups into specific regions is emphasized via confidence ellipses.

Voronoi tessellation of the two-dimensional projection plane offers an alternative to confidence ellipses [4]. Voronoi tessellation is a well-established tool in crystal structure research, where it facilitates the analysis of local atomic arrangements, grain boundaries, and phase transitions [7]. By defining neighbor relations through the identification of shared facets between Voronoi cells, this method provides a parameter-free approach to investigating local structure and nucleation processes in supercooled liquids [8]. Given these capabilities, Voronoi tessellation is well-suited to enhance the (visual) assessment of relationships among individual cases within complex group structures in diverse data sets. Its advantages include quicker visual identification of concentrations of the study group and disruptions to the expected color pattern in the Voronoi tessellation. However, further consideration is needed to determine whether Voronoi cell-based projections could replace standard confidence ellipse visualizations and included in the defaults of omics analysis software [9].

## Methods

### Confidence ellipses

Confidence ellipses illustrate the uncertainty or variability associated with group means in two dimensions. Mathematically, a confidence ellipse is a two-dimensional generalization of a confidence interval and is typically constructed under the assumption that the data within each group follows a bivariate normal distribution. For a group of projected points with a mean vector μ and a covariance matrix Σ, the confidence ellipse for a specified confidence level (e.g., 95%) is defined as the set of points satisfying the equation

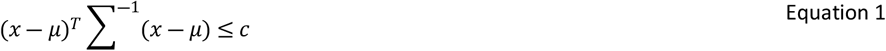

where *c* is a constant determined by the desired confidence level and related to the quantile of the chi-squared distribution with two degrees of freedom [10].

Geometrically, the confidence ellipse is centered at the group mean, and its axes are aligned with the principal directions of the covariance matrix. The lengths of the axes reflect variability in each direction. A circular ellipse indicates equal variance in both dimensions (uncorrelated variables), while an elongated ellipse indicates higher variance in one direction (correlated variables). The area inside the ellipse represents the region where the true group mean would fall with the specified probability (e.g., 95%) if the experiment were repeated many times.

### Voronoi cells

Voronoi cells [11] are defined as follows. Let {*p*_1_, *p*_2_,…,*p*_*n*_} be a set of *n* distinct points in a metric space *D* ⊆ ℝ^*d*^ with a distance function *d*(*x*,*y*) defined for all *x*,*y* ∈ *D*. The Voronoi tessellation partitions *D* into *n* regions, called Voronoi cells, each associated with a point *p*_*i*_ ∈ *P*.

The Voronoi cell *V*_*i*_ corresponding to the point *p*_*i*_ is defined as:

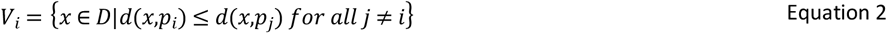

The definition of the Voronoi cell *V*_*i*_ means that for any point *x* in the domain *D*,*x* will be assigned to the cell of the point *p*_*i*_ to which it is closest, according to the distance function *d*(*x*,*y*). Formally, *x* ∈ *V*_*i*_ if and only if *V*_*i*_ = *d*(*x*,*p*_*i*_) ≤ *d*(*x*,*p*_*j*_) *for all j* ≠ *i*. This ensures that the entire space *D* is divided into non-overlapping regions, each region containing all points that are closest to a specific center *p*_*i*_.

Geometrically, the boundaries between Voronoi cells are formed by the set of points equidistant from two or more centers. In the case of standard Euclidean distance in two dimensions, for example, these boundaries are straight lines, or perpendicular bisectors, between pairs of points. In higher dimensions, the boundaries become hyperplanes. Thus, Voronoi tessellations naturally partition space based on proximity to a given set of points.

### Implementation

The visualization based on Voronoi cell tessellation of the projection plane [4] functions similarly to a “political map”. Specifically, data projections are visualized by plotting data points in a two-dimensional ℝ^2^ plane. Each point is colored according to its prior classification. Then, Voronoi cells are computed around each data point, and these cells are colored according to the same class labels. Alternatively, the coloring can reflect different classes, such as those resulting from clustering, prior classifications, or study hypotheses.

Coding was done in the R language (Ihaka and Gentleman 1996) using the R software package (R Development Core Team 2008), version 4.5.0 for Linux, with the PyCharm integrated development environment (version 2025.1.1.1; Professional Edition; JetBrains c.r.o., Prague, Czech Republic), which provides an AI Assistant plugin (https://plugins.jetbrains.com/plugin/22282-jetbrains-ai-assistant, version 251.26094.80.11), which was used on own previous code [4]. The R code uses the “deldir” R package (https://cran.r-project.org/package=deldir [12]) to compute the Voronoi tessellation of a set of two-dimensional points and assign each cell to a class. This is a novel replacement for the now-obsolete R package previously used [4], so the present code reintroduces easy generation of Voronoi tessellations for biomedical researchers. The R package “ggplot2” (https://cran.r-project.org/package=ggplot2 [13]) was used for plotting, with the colorblind-friendly palette” colorblind_pal” from the “ggthemes” package (https://cran.r-project.org/package=ggthemes [14]).

The resulting plot shows each Voronoi cell colored by class, with the original points overlaid. This allows for an intuitive visual assessment of class separation. Optionally, the cells can be colored according to an alternative classification, such as clustering. This is useful for visually judging the clustering success when contrasting it with prior classification. The code includes two functions. The more complete function, “create_projection_plots” produces three types of plots, including confidence ellipses, Voronoi tessellation, or a combination of both, whereas the reduced function, “create_voronoi_plot”, creates only the colored Voronoi cell variant that is the focus of this report. The detailed function call is described in the R code and its description provided at https://github.com/JornLotsch/voronoi_projection_plot. The function calls and parameters are shown in Textbox 1 and Table 1 respectively. An R library “VoronoiBiomedPlot” has been submitted to the Comprehensive R Archive Network (CRAN) (initial check pending).

**Table 1:** Function call parameters to invoke various Voronoi tessellations and/or confidence ellipse-style plots of two-dimensional data with (or without) class structure, and list of main outputs of the function. Compare https://github.com/JornLotsch/voronoi_projection_plot.

| Parameter/Output | Type | Default | Description |
| --- | --- | --- | --- |
| **Function call parameters** |  |  |  |
| **Data** | data.frame | - | Data with $\geq 2$ numeric columns for coordinates |
| **class_column** | character/vector | NULL | Column name or vector of class labels |
| **alternative_class_column** | character/vector | NULL | Alternative column name or vector of class labels |
| **coordinate_columns** | character vector | NULL | Column names to use as coordinates (if NULL, uses first 2 numeric columns) |
| **case_labels** | character vector | NULL | Individual case labels (uses row numbers if NULL) |
| **coord_names** | character vector | c("Dim1", "Dim2") | Names for coordinate axes |
| **Title** | character | NULL | Plot title |
| **show_labels** | logical | FALSE | Whether to show case labels |
| **voronoi_alpha** | numeric | 0.3 | Transparency of Voronoi regions (0-1) |
| **point_size** | numeric | 2 | Size of data points |
| **legend_position** | character/numeric | "bottom" | Legend position |
| **color_palette** | function/character | NULL | Custom color palette |
| **add_grid_lines** | logical | FALSE | Whether to add origin grid lines |
| **color_points** | character | "primary" | Which classification to use for point colors ("primary" or "alternative") |
| **fill_voronoi** | character | "primary" | Which classification to use for Voronoi fills ("primary" or "alternative") |
| **point_shape** | character | "none" | Which classification to use for Voronoi fills ("primary", "alternative" or "none") |
| **label_fontface** | character | "plain" | Font face for case labels ("plain", "bold", "italic", "bold.italic") |
| **label_size** | numeric | 3.88 | Size of case labels |
| **Outputs** |  |  |  |
| **result\$scatter_plot** | ggplot | | Standard scatter plot of the projected data |
| **result\$voronoi_plot** | ggplot | | Voronoi tessellation plot with data points |
| **result\$combined_plot** | ggplot | | Combined visualization with additional features |

### Data sets

#### Artificial data

##### Two-class synthetic benchmark data set

The synthetic dataset comprises nine variables acquired from n = 80 cases from two different classes, *C*_1_(*n*_1_ = 40) and C_2_ (n_2_=40). Each observation is represented as a vector in ℝ^9^, corresponding to nine continuous variables with varying degrees of class separation. Variables *A* and *B* are sampled from normal distributions with class-specific parameters. For variable *A*, the distribution within class *C*_*i*_ is 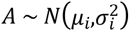, where (*µ*_1_,*µ*_2_) = (4,8) and (*σ*_1_,*σ*_2_) = (1,1). Variable *B* is generated using scaled and cross-referenced parameters, i.e., 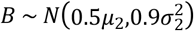 for class *C*_2_ and 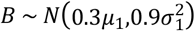 for class *C*_1_, creating an inverse relationship between classes. Variables *E* and *G* are drawn from uniform distributions *E* ∼ *U*(*a*_*i*_,*b*_*i*_) for each class *C*_*i*_, where the bounds are derived from jittered class-specific mean values to ensure class separation. Variables *C* and *D* are constructed via weighted sampling from sequential values and subsequent jittering, introducing moderate variability with partial class overlap. Variables *H* and *I* are sampled from a common uniform distribution across all classes, with variable *I* being a jittered linear transformation of *H*. Data set generation details are available at https://github.com/JornLotsch/voronoi_projection_plot/blob/main/Two_class_artifical_data_example.R.

An alternative version of this dataset was created in which the class labels of three cases in each group were reassigned to the opposite class. This modification introduced controlled misclassification, enabling the evaluation of visualization methods under conditions of label noise and their ability to identify atypical or mislabeled instances within well-separated groups.

##### Three-class synthetic data sets in a significant and a non-significant variant

The second synthetic data set also comprises nine variables, this time acquired from n = 75 cases from three different classes, *C*_1_(*n*_1_ = 20), *C*_2_(*n*_2_ = 40) and *C*_3_(*n*_3_ = 15). Each observation is again represented as a vector in ℝ^9^. Variables *A* and *B* are sampled from normal distributions with class-specific parameters. For variable *A*, the distribution within class *C*_*i*_ is 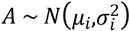, where (*µ*_1_,*µ*_2_,*µ*_3_) = (4,6,8) and (*σ*_1_,*σ*_2_,*σ*_3_) = (2,4,3). Variable *B* is generated analogously, using scaled versions of the same parameters. Variables *E* and *G* are drawn from uniform distributions *E* ∼ *U*(*a*_*i*_,*b*_*i*_) for each class *C*_*i*_. Variables *C* and *D* are constructed via weighted and jittered sampling from uniform distributions, introducing moderate variability and partial class overlap. Variables *H* and *I* are sampled from a common uniform distribution across all classes with minimal class discrimination. In the present random realization of the data set, group differences happened to be significant at p < 0.05, according to Kruskal-Wallis tests [15], for variables *A, B, E* and *G*. Data set generation details are available at https://github.com/JornLotsch/voronoi_projection_plot/blob/main/Three_class_artifical_data_example.R.

An alternative version of the above data set without significant group differences (all Kruskal-Wallis tests: p > 0.05) was generated by independently permuting each variable in the original dataset, thus destroying the association between the variables and class membership.

##### Clustering problem data set

The data set used to illustrate the limitations of clustering algorithms such as k-means was taken from the Fundamental Clustering Problems Suite (FCPS), a benchmark collection designed to expose common challenges encountered in real-world clustering tasks [16]. The FCPS suite includes artificial datasets constructed to highlight specific structural difficulties for clustering algorithms, such as varying cluster shapes, densities, and separations. Selected for this study, the two-dimensional “Lsun” dataset comprises 400 samples distributed across three well-separated classes. These classes differ markedly in their geometric properties: one forms a roughly spherical cluster, while the other two are rectangular (“brick-shaped”) and vary in size. This configuration introduces varying cluster variances and densities, presenting a significant challenge for algorithms like k-means [17, 18], which assumes that clusters are hyperspheric and therefore perform poorly on the “Lsun” dataset [19]. The clustering demonstration was done using the “kmeans” function from the R “stats” core package, setting the parameters centers = 3 (the original class number in the “Lsun” dataset) and nstart = 100; for details, see the R code at https://github.com/JornLotsch/voronoi_projection_plot/blob/main/Lsun_example.R.

#### Biomedical data

##### Sensitivity to experimental nociceptive stimuli (“QSTpainEJPtransf”)

This psychophysical pain-related dataset (“QSTpainEJP”) was available from an in-house study on clinical quantitative sensory testing (QST) involving 127 healthy subjects [20, 21]. The data set includes 22 sensory measures of pain, i.e., nine from classical pain models (e.g. pressure, cold, electric, chemical and laser-evoked pain thresholds and intensities), and ten from a clinically established QST battery [22]. The QST parameters encompass a range of thermal and mechanical pain and sensation thresholds. Measures were harmonized to ensure that higher values indicate greater pain and were log-transformed as appropriate. Detailed protocols and variable descriptions are available in the original publications [20, 21]. The data set is publicly accessible via the Mendeley Data repository (DOI: 10.17632/9v8ndhctvz.1) at https://data.mendeley.com/preview/9v8ndhctvz. For the present purpose of data visualization, only the n = 72 complete cases were projected to avoid imputation of missing’s (34 men and 38 women). Since one of the previously reported results had been a sex difference in pain thresholds [23], sex was used as the data set’s class structure for the present purpose (coding: 1 = male, 2 = female). For details, see the R code at https://github.com/JornLotsch/voronoi_projection_plot/blob/main/pain_data_example.R.

##### Clinical lipidomics of rheumatic diseases (“PsA_lipidomics”)

This lipidomics dataset (“PsA_lipidomics”) includes plasma lipid profiles from 81 patients diagnosed with psoriatic arthritis (PsA) and 26 healthy control subjects matched by age and sex. This results in a data matrix of 107 subjects and 292 lipid markers. Samples were collected as part of a cross-sectional study at a university-based tertiary care rheumatology center. Both targeted and untargeted assays were employed to capture a broad spectrum of lipid classes, including carnitines, ceramides, glycerophospholipids, sphingolipids, glycerolipids, fatty acids, sterols, esters, and endocannabinoids.

During analysis, a subset of control samples was observed to cluster anomalously with PsA patients. Further investigation revealed that these outliers were due to batch effects; control samples processed in separate batches exhibited divergent data projections, indicating insufficient normalization and batch correction. These characteristics made the dataset suitable for comparing standard confidence ellipse plots with the proposed Voronoi tessellation approach, with a focus on identifying and interpreting problematic or mislabeled cases. The dataset is publicly available via the Mendeley Data repository (DOI: 10.17632/32xts2zxdc.1) at https://data.mendeley.com/datasets/9v8ndhctvz/1, where additional methodological details can be found. Its processing for this report was similar to that of the above pain-related data set (https://github.com/JornLotsch/voronoi_projection_plot/blob/main/PsA_data_example.R).

##### COVID-19 metabolomics data set (“covid_metabolomics”)

Another example was processed using the MetaboAnalyst 6.0 web-based platform [9], which is a comprehensive tool for metabolomics data analysis available at https://www.metaboanalyst.ca/. Specifically, an online statistical analysis was performed at https://dev.metaboanalyst.ca/MetaboAnalyst/upload/MultifacUploadView.xhtml on liquid chromatography-mass spectrometry (LC-MS) peak intensity data from 59 samples, including 20 healthy controls and 39 patients with SARS-CoV-2 infection (data from https://api2.xialab.ca/api/download/metaboanalyst/covid_metabolomics_data.csv, metadata from https://api2.xialab.ca/api/download/metaboanalyst/covid_metadata_multiclass.csv).

This resulted in a data matrix of 59 samples by 2,054 peaks (m/z and retention time). Four metadata factors were included: Diagnosis, Gender, Treatment, and Age. Data preprocessing involved variance filtering using a nonparametric relative standard deviation (median absolute deviation normalized by median), abundance filtering based on median intensity values, median normalization, and autoscaling (mean-centering followed by division by the standard deviation of each variable). The platform’s interactive PCA was conducted, and the resulting loading data was downloaded and plotted for the present analysis.

## Results

### Visualization of clear class structures

In data sets with two classes that are clearly separated on the ℝ^2^ plane of a PLS-DA projection (Figure 1), visualization using confidence ellipses is adequate (Figure 1 A). Notably, the Voronoi tessellation provides an equally clear alternative (Figure 1 B), supporting its suitability as a visualization method comparable to the current standard. However, when three points per class are switched to the respective opposite class, the irregularity may escape attention in the ellipses plot (Figure 1 C), but it is clearly evident in the Voronoi tessellated plot (Figure 1 D).

**Figure 1:**
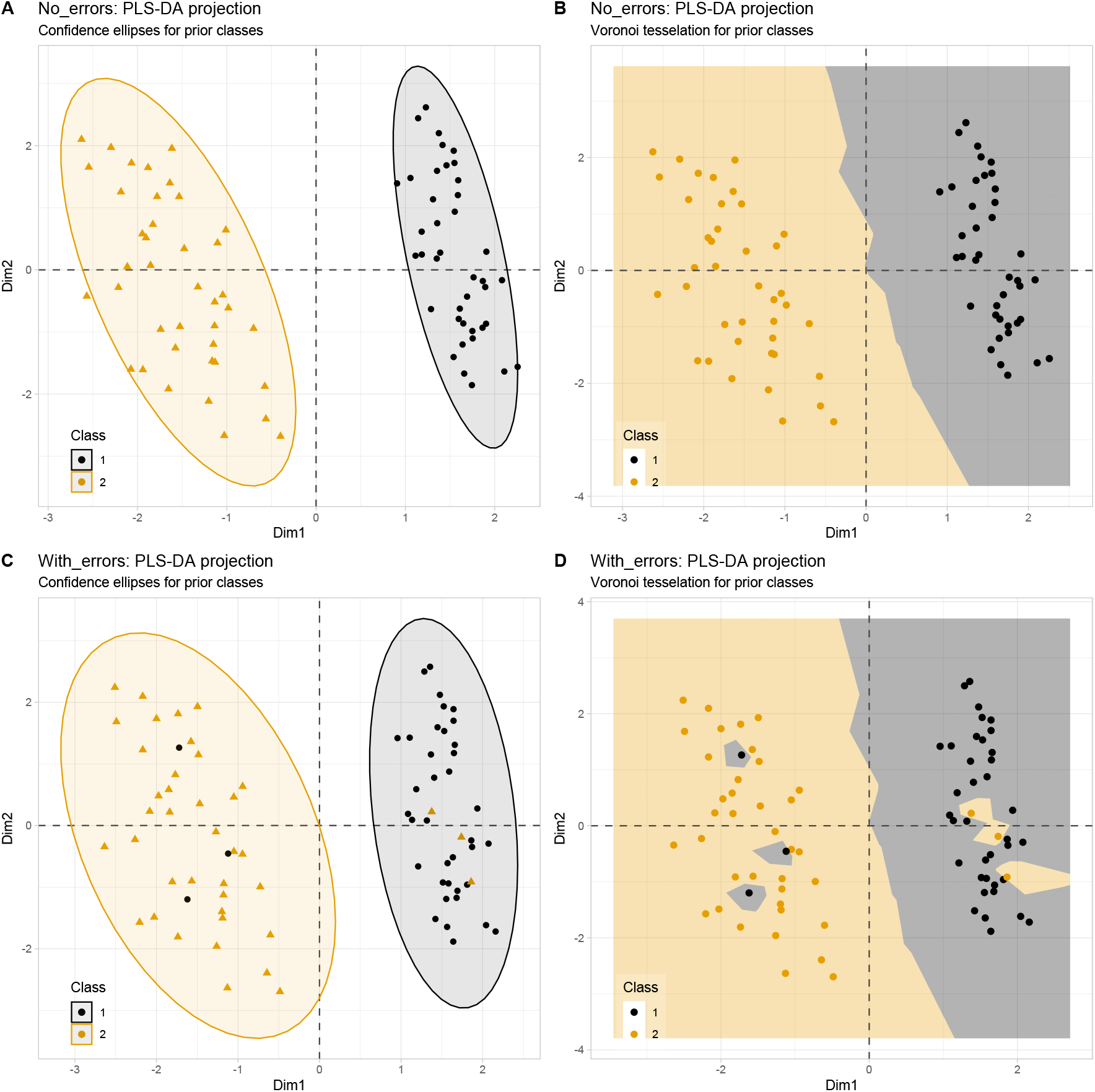
PLS-DA projection results for a nine-dimensional, two-class synthetic benchmark dataset (*n*_1,2_= 40/40). **A:** Standard 95% confidence ellipse visualization on the ℝ^2^ plane with color coding for the original (prior) classes. **B:** The same projection as in panel A, but with Voronoi tessellation instead of confidence ellipses. **C and D:** Introduction of class aberrations: Three instances per class were assigned to the opposite class, after which the PLS-DA projection was repeated. This resulted in only minor changes to the projection. However, panels C and D demonstrate the visualization of the aberrant cases, which are almost hidden in C and very clear in D.

### Performance in complex and ambiguous separation scenarios

In more complex scenarios where three groups differ to varying degrees across several variables, both Voronoi tessellation and confidence ellipses provide consistent visualizations (Figure 2 A and B). The tessellated plot (Figure 2 B) may reveal slightly more pronounced group segregation, but it does not clearly outperform the confidence ellipse plot in clarity. However, when the dataset is permuted, confidence ellipses become less decisive in indicating the now absent significant group segregation (Figure 2 C and D). This leaves both segregated and non-segregated group structures as possible interpretations (Figure 2 C). In contrast, Voronoi tessellation clearly shows the lack of meaningful group segregation in the permuted data. The cells disperse across the ℝ^2^ plane without any evident concentration or pattern (Figure 2 D). Thus, Voronoi tessellation appears superior in distinctly conveying the absence of group structure in ambiguous cases.

**Figure 2:**
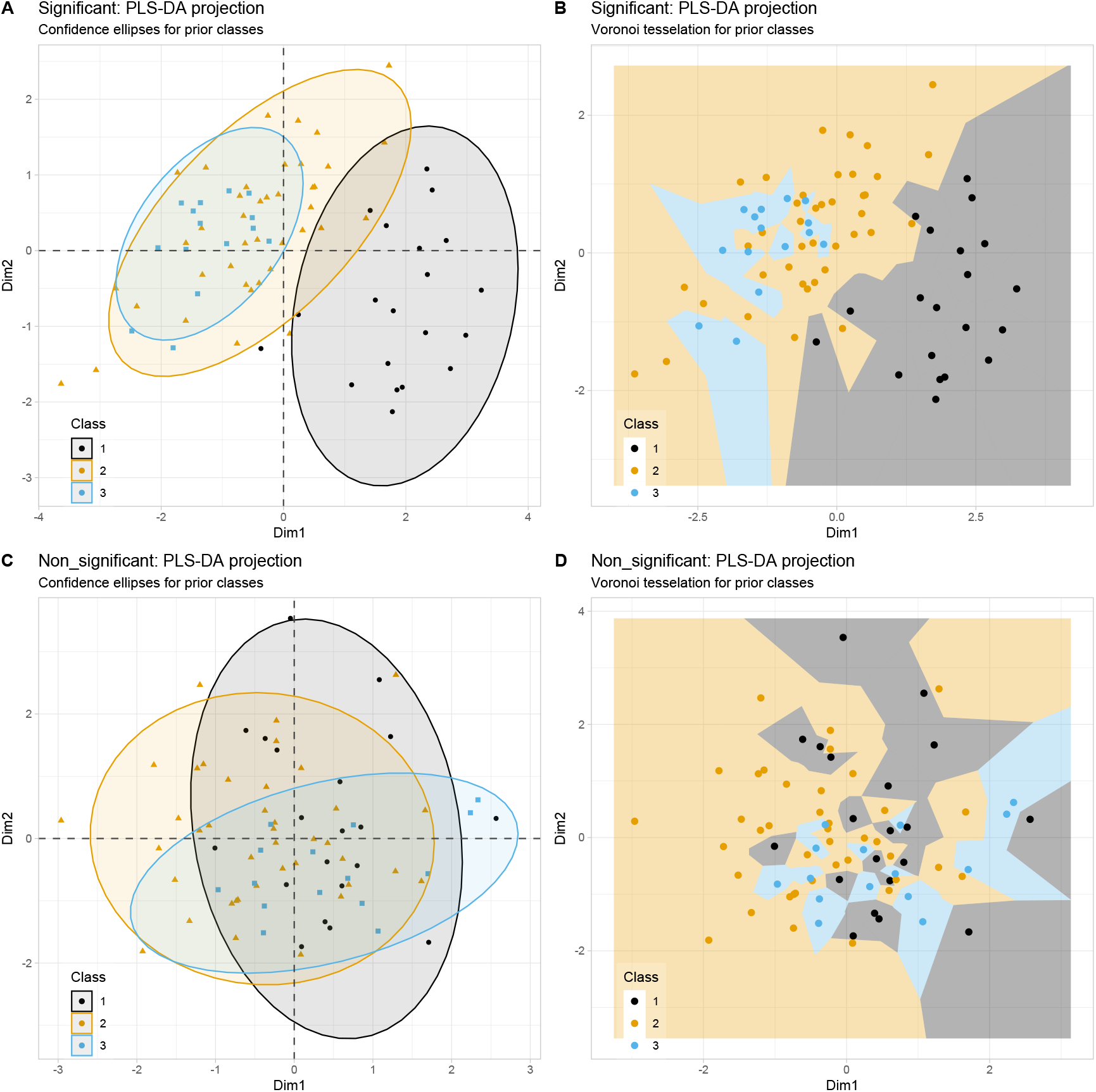
PLS-DA projection for a nine-dimensional, three-class synthetic dataset (*n*_1,2,3_ = 20/40/15) with significant and non-significant variants. **A:** Standard 95% confidence ellipse visualization on the ℝ^2^ plane, with color coding for the original classes in the significant data scenario. **B:** The same projection as in panel A, but with Voronoi tessellation instead of confidence ellipses. **C** and **D:** Results of PLS-DA projection following randomly permuting each variable to make the class differences non-significant.

### Detection of “internal outliers” in biomedical data

A more detailed visualization of group structure is achieved using Voronoi tessellation in the biomedical, pain-related “QSTpainEJP” data set (Figure 3). The separation of groups defined by sex is evident in both displays. Standard outliers, such as cases #1356, 1365, 1375, 1382, and 1474, can be easily identified when case labeling is enabled (see Textbox 1), as they are significantly distant from the primary clusters. In the Voronoi tessellation, additional cases (#1299, 1378, and 1451) are scattered among the opposite sex and are highlighted by their Voronoi cells that border those of the other group. These “internal outliers” may indicate data or laboratory errors, novel phenotypes, or unconsidered conditions, necessitating further investigation.

**Figure 3:**
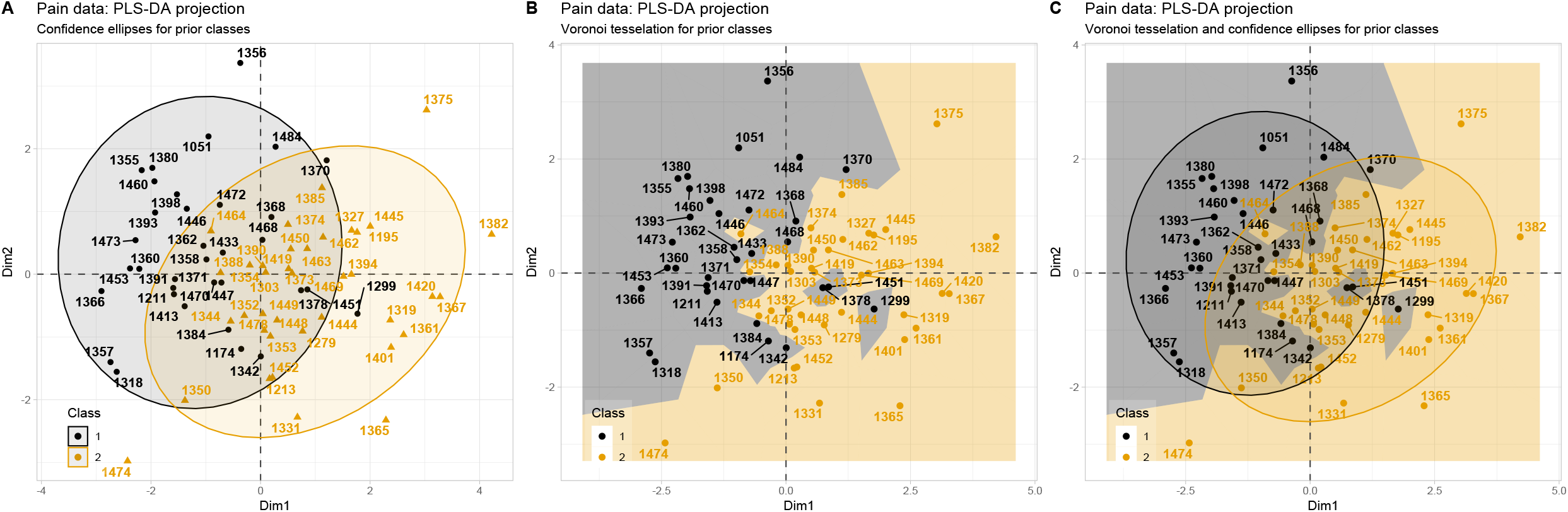
PLS-DA projection results for quantitative sensory testing (QST) pain data from 72 healthy subjects (34 men, 38 women) across 19 pain-related variables (dataset “QSTpainEJP”). Individual cases are labeled to enable follow-up analysis at the single-case level. **A:** Standard confidence ellipses on the ℝ^2^ plane with color coding for sex groups. **B:** shows the same projection as panel A, but with Voronoi tessellation instead of confidence ellipses. **C:** Combined visualization of both approaches.

A similar pattern emerged in the analysis of the PsA_lipidomics dataset, where PLS-DA projections revealed irregularities in the group structure due to batch effects (Figure 4). Notably, several PsA patient samples were scattered among the healthy controls, while most patients clustered together as expected. The Voronoi tessellation more clearly highlighted this dispersion than confidence ellipse visualizations did. Controls such as #K112, K111, and K120 projected at the periphery of their group or in proximity to patient samples and corresponded to cases with known batch normalization issues. Thus, the Voronoi approach provided enhanced sensitivity for detecting problematic or mislabeled cases by emphasizing group boundaries and internal outliers more effectively than standard methods.

**Figure 4:**
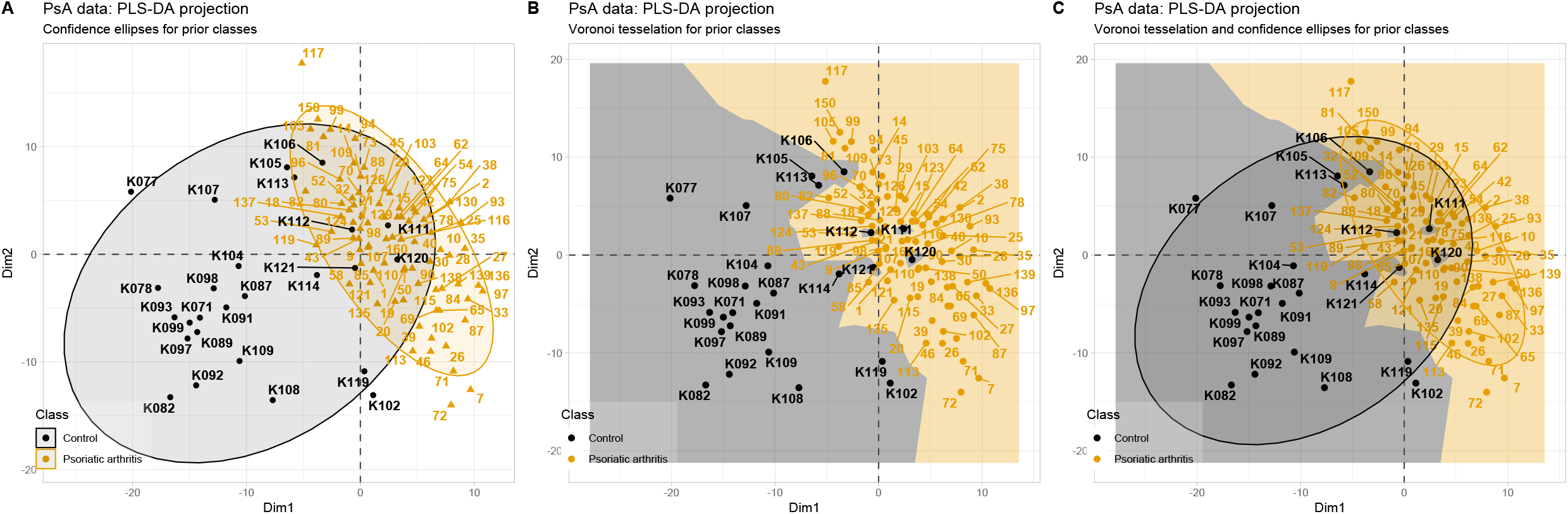
PLS-DA projection results for lipidomic data from 81 patients with psoriatic arthritis (PsA) and 26 healthy controls across 292 lipid markers (dataset “PsA_lipidomics”). Individual cases are labeled to enable follow-up analysis at the single-case level. **A:** Standard confidence ellipses on the ℝ^2^ plane with color coding for diagnostic groups. **B:** The same projection as panel A, but with Voronoi tessellation. The controls (ID code starting with “K”) projected among PsA patients (grey spots within a yellow zone) are among cases with failed laboratory batch correction. **C:** Combined visualization of both approaches.

The “covid_metabolomics” dataset further demonstrated the advantages of Voronoi tessellation over standard visualization techniques for analyzing the effects of SARS-CoV-2 on metabolism (Figure 5). Although group separation between patients with and without the disease was apparent in the PCA projection, the Voronoi tessellation provided clearer identification of outliers and cases whose projections overlapped with the opposite class. These observations are consistent with those from other biomedical datasets. Please also compare the default plot style from the MetaboAnalyst platform in Figure 5 D. It should also be noted that the analysis of this dataset was primarily conducted to create a plot suitable for the current presentation. To generate scientifically valid results on the metabolomic pattern of patients with COVD-19, the data processing steps and resulting projections may differ.

**Figure 5:**
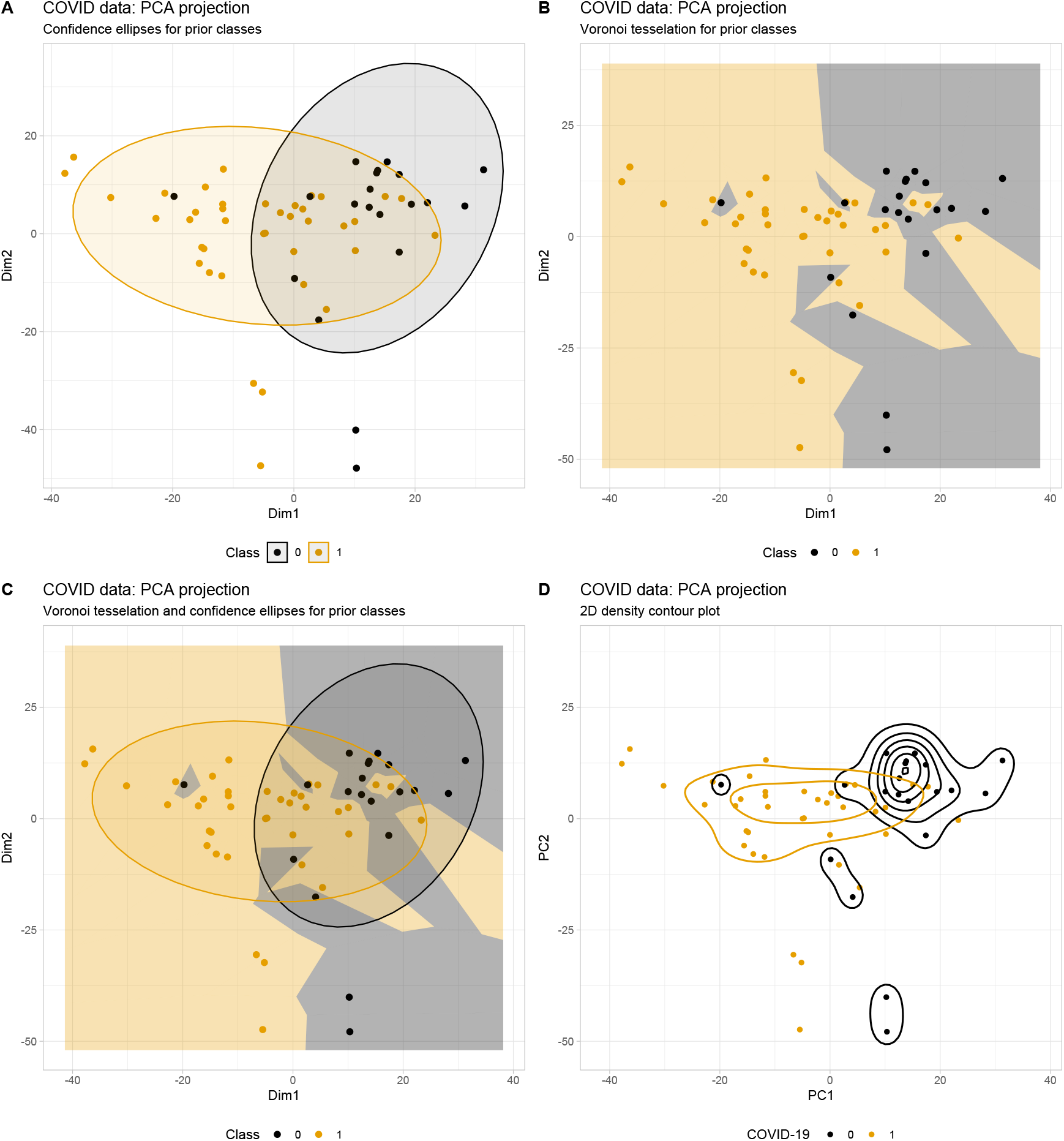
PCA projection results for metabolomics data from 39 COVID-19 patients and 20 healthy controls across metabolite markers. Individual cases are labeled to enable follow-up analysis at the single-case level. The data processing and PCA had been performed directly at the MetaboAnalyst 6.0 web site at https://www.metaboanalyst.ca/. **A:** Standard confidence ellipses on the ℝ^2^ plane with color coding for diagnostic groups. **B:** The same projection as panel A, but with Voronoi tessellation. The tessellation reveals clear spatial separation between COVID-19 patients and healthy controls, however, with overlap between groups. **C:** Combined visualization of both approaches. **D:** Dot plot overlaid with a 2D density contour plot analogous to the MetaboAnalyst 6.0 [9] default.

### Class structure detection and agreement analysis

Voronoi tessellation also proved useful for visualizing class or cluster structures and highlighting clustering failures. For example, the “Lsun” dataset, constructed with two rectangular clusters, typically defies correct clustering by the k-means algorithm, as previously shown [16, 19]. The Voronoi tessellation plot improves visibility of the original class structure (Figure 6 A). As expected, k-means clustering failed to capture this structure, instead producing clusters that corresponded better to its inherent assumption of circular shapes in this two-dimensional dataset. By switching the coloring of data points and Voronoi cells between true classes and clusters using different combinations of the function parameters (“class_column”, “alternative_class_column”, “color_points”, “fill_voronoi”; see Textbox 1), the incorrect clustering became clearly visible (Figure 6B and C).

**Figure 6:**
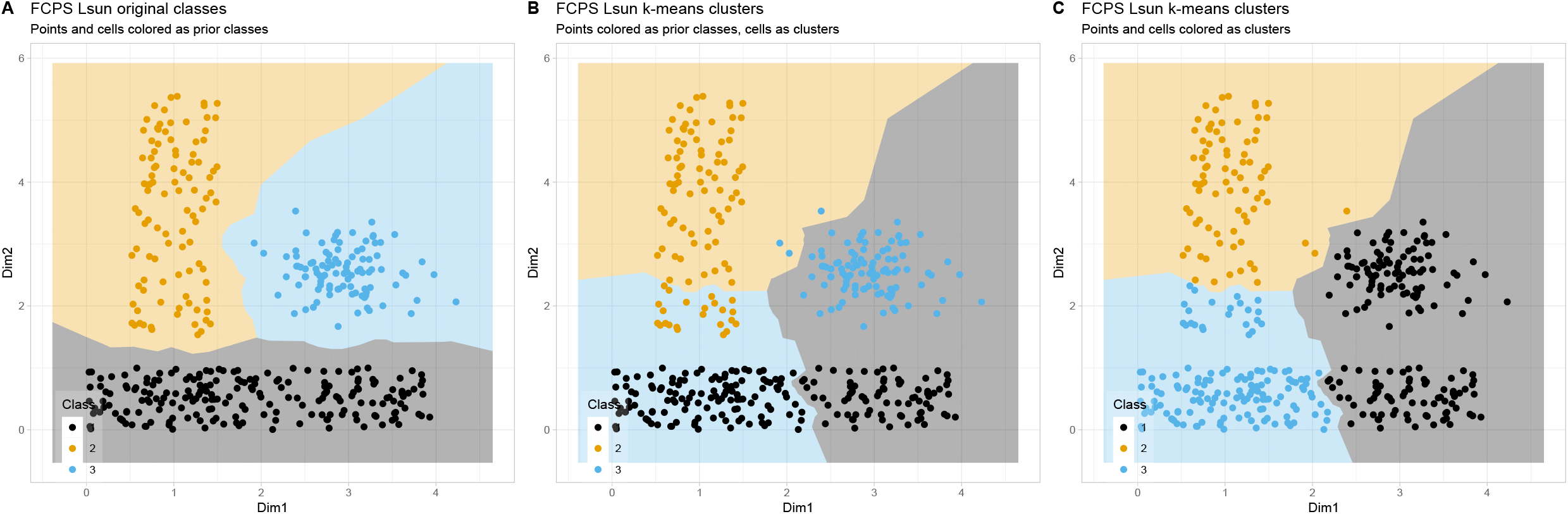
Voronoi tessellation of a 2d plot of the original (non-projected) FCPS [16] artificial “Lsun” dataset comparing original class structure with k-means clustering results (k=3). **A:** Original class structure, with points and cells colored according to prior classes. **B:** Comparative visualization in which the points are colored according to the original classes and the Voronoi cells are colored according to the k-means cluster assignments, showing the agreement and disagreement between the prior classification and the unsupervised clustering. **C:** k-means clustering results, with both points and Voronoi cells colored according to cluster assignments.

## Discussion

The strength of Voronoi tessellation in data projection planes lies in its ability to highlight disruptions in expected patterns. This qualifies it as the method of choice in crystallography [7, 8], and as the current demonstrations indicate, it appears to qualify for omics research as well. The tessellation of the projection plane makes it easier to identify data points that deviate from their fellow class members with respect to the main projection direction. The visualization effectively emphasizes what might be termed “internal outliers”. Conversely, conventional (“external”) outliers may be less visually prominent in this framework, though they can still be spotted easily, although the definition of outliers can vary [24]. However, combining Voronoi tessellations with confidence ellipses may be a complementary visualization strategy offering statistical and geometric perspectives on data clustering. Confidence ellipses reveal distributional characteristics. Voronoi diagrams, on the other hand, provide an intuitive representation of decision boundaries based on nearest-neighbor proximity. This dual approach leverages the strengths of both methods to offer a more comprehensive view of group structure and enhanced outlier detection capabilities.

Voronoi diagrams divide space into regions based on proximity to a set of points, which gives a clear visual partition of influence or dominance regions, which is ideal for spatial or clustered data, such as ecological zones, cellular territories, or geographical mapping [25]. Voronoi cells inherently define local neighborhoods based on distance, unlike PCA or PLS-DA plots, which may distort neighborhood relationships due to projection into lower-dimensional space, this is especially helpful in data where local context matters, like tissue analysis in histology or population distribution. [26, 27]. Despite that, PCA and PLS-DA compress high-dimensional data into 2 or 3 dimensions, which can lead to loss of important variance or discriminative features. Voronoi tessellation can work directly with 2D spatial data without such compression, preserving more truthful geometrical relationships [28-30].

Incorporating Voronoi tessellation is an efficient approach to visualizing class boundaries in projected data spaces. However, it is not limited to projected data, as demonstrated by the “Lsun” data set example where clustering was done on the non-projected data. Unlike traditional clustering visualizations, which rely solely on point distributions, Voronoi regions provide explicit territorial boundaries for each observation. This creates an intuitive classification map of the feature space, which facilitates identifying potential misclassification zones, interpreting the local neighborhood structure around individual data points, and detecting regions of class overlap that may not be apparent in conventional scatter plots. That is, the visualization provides a diagnostic signal that points to data set instances that deviate in their projection from the expectations based on group membership. Whether these deviations are due to laboratory or data errors, represent a novel finding, or merely express normal data variation, is then subject to interpretation by the research team. The visualization merely helps identify cases for investigation. There, Voronoi tessellations are especially useful in semi-supervised settings where they enable comparisons between data projections and prior classifications. This includes assessing whether the projected data supports existing classifications. However, the visualization is not limited to semi-supervised settings. In unsupervised settings, discrepancies between clustering results and the underlying structure can be easily identified through visual inspection. For instance, as illustrated in the “Lsun” example (Figure 6C), a solitary black cluster on the right appears as two dense regions upon examination using Voronoi tessellation. This highlights shortcomings in the clustering approach.

### Strengths and Limitations

In the actual implementation, support for dual classification systems (primary and alternative; see Textbox 1) addresses a common challenge in multivariate analysis, where data points may belong to multiple taxonomic or functional categories simultaneously. This flexibility enables researchers to explore different classification schemes within the same projected space and compare how different grouping strategies affect visual separation patterns. Additionally, limited availability of software coded solutions highlights the need for an easy-to-use implementation to facilitate the application of this data projection and/or clustering visualization method.

While this visualization approach provides insights into the class structure of a data set, the effectiveness of the method depends on the quality of the initial dimensionality reduction. Poor projection techniques may obscure rather than reveal class relationships [31]. Therefore, choosing the right projection techniques remains a key component of the data analysis workflow. Although each data set is unique, one can rely on available comparative benchmarking results [4, 19], although *a priori* definition of the best projection method, including “none”, is difficult. Additionally, the Voronoi tessellation approach assumes that proximity in the reduced space corresponds to similarity in the original, high-dimensional space, which is not always the case. Future developments could incorporate uncertainty measures for dimensionality reduction and explore alternative distance metrics for tessellation.

## Conclusions

This study introduces a visualization framework integrating Voronoi tessellation and optional confidence ellipses to improve the evaluation of data projections and clustering results. Across artificial and biomedical datasets, the method demonstrated superior sensitivity in detecting structural inconsistencies between projected data and known classifications. It outperformed conventional scatter plots with confidence ellipses alone. Notably, the method enabled clearer identification of cases in which the clustering results did not align with the underlying group structures. These findings establish Voronoi tessellation as a robust alternative to standard confidence ellipse visualizations in omics analysis software and highlight the added value of a combined visualization strategy. Overall, our approach provides a practical and effective tool for more reliably interpreting dimensionality reduction and clustering in various data analysis applications.

## Declarations

### Ethics approval

Not applicable. All datasets used in this report were obtained from publicly accessible repositories, which are specifically referenced alongside the datasets.

### Consent for publication

Not applicable. All datasets used in this report were obtained from publicly accessible repositories, which are specifically referenced alongside the datasets.

### Availability of data and material - (data transparency)

The data used in the experiments in this report are publicly available and referenced accordingly. They can be accessed at https://github.com/JornLotsch/voronoi_projection_plot (generation code for the first two artificial data sets), at https://doi.org/10.3390/data5010013 (FCPS “Lsun” data set), at https://data.mendeley.com/datasets/9v8ndhctvz/1 (pain related data set) and at https://data.mendeley.com/datasets/32xts2zxdc/1 (PsA lipidomics data set).

### Code availability - (software application or custom code)

Relevant R code is available at https://github.com/JornLotsch/voronoi_projection_plot. Upload of an R library “VoronoiBiomedPlot” on the Comprehensive R Archive Network (CRAN) is pending.

### Authors’ contributions

JL – Conceptualization of the project, developing the project, theoretical background, programming, performing experiments, writing of the manuscript, data analyses and creation of the figures, funding acquisition.

DK – Evaluation of the feasibility of the approach for biomedical researchers, contributing to the writing of the manuscript.

## Acknowledgements

Not applicable.

---

**Textbox 1:**
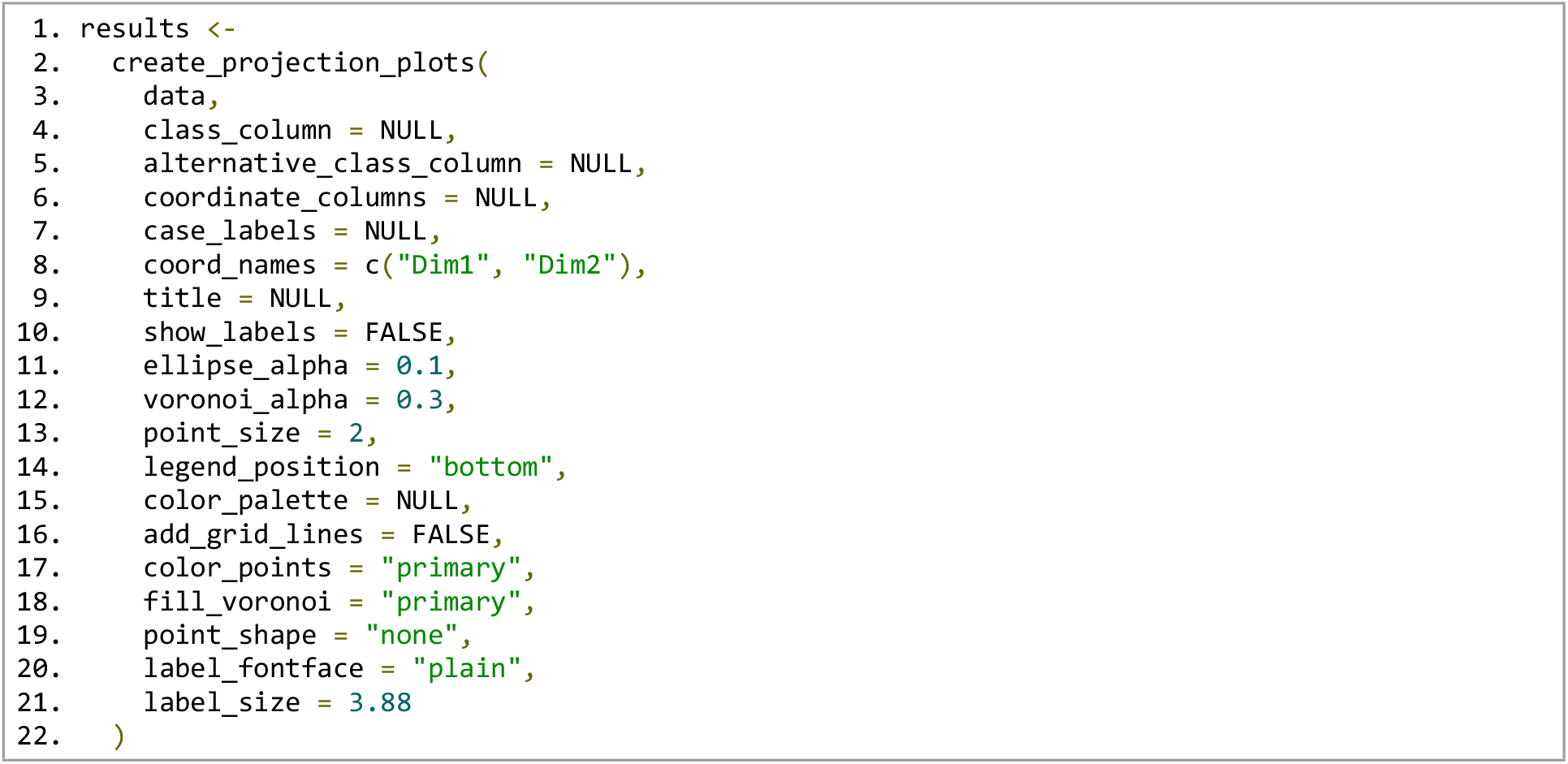
R function calls for the 2D visualization of the Voronoi-tessellated data projection. For parameter definitions, see Table 1 and https://github.com/JornLotsch/voronoi_projection_plot.

